# Altered cortical gyrification in adults who were born very preterm and its associations with cognition and mental health

**DOI:** 10.1101/2019.12.13.871558

**Authors:** Chiara Papini, Lena Palaniyappan, Jasmin Kroll, Sean Froudist-Walsh, Robin M Murray, Chiara Nosarti

## Abstract

**Background:** The last trimester of pregnancy is a critical period for the establishment of cortical gyrification and altered folding patterns have been reported following very preterm birth (< 33 weeks of gestation) in childhood and adolescence. However, research is scant on the persistence of such alterations in adulthood and their associations with cognitive and psychiatric outcomes.

**Methods:** We studied 79 very-preterm adults and 81 age-matched full-term controls. T1-weighted images were used to measure local gyrification index (LGI), indicating the degree of folding across multiple vertices of the reconstructed cortical surface. Group and group-sex LGI differences were assessed using per-vertex adjustment for cortical thickness and overall intracranial volume. Within-group correlations were also computed between LGI and functional outcomes, including general intelligence (IQ) and psychopathology.

**Results:** Very preterm adults had significantly reduced LGI in extensive cortical regions encompassing the frontal, anterior temporal and occipito-parietal lobes. Alterations in lateral fronto-temporal-parietal and medial occipito-parietal regions were present in both males and females, although males reported more extensive alterations. In both very preterm adults and controls, higher LGI was associated with higher IQ and lower psychopathology scores, with the spatial distribution of these associations substantially differing between the two groups.

**Conclusions:** Very preterm adults’ brains are characterized by significant and widespread local hypogyria and these abnormalities might be implicated in cognitive and psychiatric outcomes. Gyrification reflects an early developmental process and provides a fingerprint for very preterm birth.

## Introduction

Preterm birth (< 37 weeks of gestation) represents over 10% of live births worldwide (1) and is associated with long-term morbidities that include psychiatric and cognitive problems (2, 3). The third trimester of pregnancy, when most preterm babies are born, is a critical period for brain development not only because of the dramatic increase of cortical surface and brain volume (4–6), but also for the establishment of the gyri and sulci (7). Gyrification, the folding of the cortical surface, occurs mainly in fetal life (8): the primary fissures (e.g. interhemispheric fissure) become apparent from 10-14 weeks of gestation and the sulcogyral morphology resembles the adult brain by the end of pregnancy (7). The overall ratio between buried and unburied cortex reaches adult complexity in the early postnatal period and remains relatively constant throughout development (8–11), with relatively smaller changes occurring in the first years of life (12) and beyond (13–15).

Therefore, preterm babies are at higher risk of irreversible alterations in cortical gyrification. Although sulci appear at a similar rate in babies born preterm and typically developing foetuses, the transition to the extra-uterine environment is a disruptive event for the gyrification process (16). Preterm newborns show typical but delayed gyral development at term equivalent age (17, 18) and exhibit less complex convolution when compared to term-born controls (19, 20). Moreover, the degree of gyrification increases with gestational age (17, 21, 22).

Altered patterns of gyrification following very preterm birth have been reported in childhood and adolescence. Compared to term-born controls, very preterm children showed a 4% decrease of global convolution in both hemispheres, as well as regional patterns of increased gyrification (hypergyria) in the medial frontoparietal cortex and decreased gyrification (hypogyria) in the temporal cortex (23). However, mixed evidence of increased gyrification (24) and reduced sulcal surface area (25) in the temporal lobe have also been reported among preterm children. Furthermore, studies investigating the morphology of the orbitofrontal cortex in adolescence found reduced depth in the secondary sulci (26), but also wider sulcagyral abnormalities (27). Regional measures of gyrification in the left temporal and prefrontal cortex have been associated with language development (24) and parent-reported executive functions (27) respectively, but not with psychiatric outcomes (27). Overall, aberrant cortical folding configurations among preterm individuals appear to persist into adolescence and may be implicated in diverse and long-lasting neurobehavioral outcomes. Alterations in cortical gyrification in later life and their implications for cognitive and psychiatric outcomes are yet to be fully understood.

Regarding adult life, Hedderich *et al*. (28) recently investigated mean cortical curvature (29) in adults born preterm and reported extensive abnormalities across the cortical mantle, with higher magnitude of deviations associated with lower gestational age, lower birth weight and more severe medical complications at birth. This study also found associations between mean curvature and intelligence, but it did not include any psychiatric measure. Recent cross-sectional studies have revealed aberrant cortical folding in psychiatric disorders such as schizophrenia (30), bipolar disorder (31), as well as depression (32) and anxiety (33). Higher rates of mood, anxiety and psychotic disorders have been reported among preterm (3) and low birth weight individuals (34). Using the Comprehensive Assessment of At Risk Mental States (CAARMS) (35) with a dimensional approach, we recently demonstrated that adults who were born preterm experience a significant psychopathological burden and exhibit a non-specific psychiatric risk associated with lower IQ (36). Such low subclinical specificity might lead to an under-recognition of mental health problems that fall below standard diagnostic thresholds. Since cortical folding is considered a powerful indicator of early neurodevelopmental pathology (37), the identification of gyrification biomarkers of psychopathological risk following preterm birth would enable to recognize and monitor individuals who are vulnerable to develop both clinical and sub-clinical psychopathology.

In this study, we assessed a cohort of very preterm adults and a group of age-matched controls who received extensive MRI, neurocognitive and psychiatric assessment, providing measures of gyrification, intelligence and mental health. Specifically, to quantify cortical gyrification, we used local gyrification index (LGI) (38), a 3D version of the gold-standard gyrification index used in post-mortem studies (11) that reflects the amount of cortex buried underneath the surface of spherical cortical mantle. LGI is also relatively independent of the thickness of the cortical mantle, unlike the mean curvature (39). The aims of this study are twofold: first, to extend previous findings (28) by using LGI to assess gyrification differences across the whole brain between adults who were born very preterm and term-born individuals and relate this to general intelligence; second, to investigate whether regional variations in gyrification relate to mental health outcomes among preterm born adults. Based on previous findings, we hypothesized that preterm born participants would present different gyrification patterns, mainly hypogyria (26, 28), but also hypergyria (23, 25), compared to controls. Furthermore, we hypothesized positive correlations between LGI and general intelligence for very preterm individuals (28) and for controls (40). Finally, we explored the associations between LGI and psychiatric symptoms in cases and controls.

## Methods and materials

### Study population

Preterm participants were recruited from a cohort of infants who were born at less than 33 weeks of gestation between 1979 and 1984 and admitted to the neonatal unit of University College Hospital, London, UK. After discharge from hospital, survivors were enrolled in a longitudinal study that involved repeated assessments in childhood (41, 42), adolescence (43) and adulthood (36, 44). Retention rate over time has been reported elsewhere (36, 45). Of the 152 subjects recruited for the most recent follow-up (at age 30), here we studied a sample of 83 very preterm adults.

Eighty-three aged-matched control participants were recruited from advertisements in the local community. Inclusion criteria were full-term birth (38-42 weeks of gestation) and birth weight > 2,500 grams. Exclusion criteria were birth complications (e.g. endotracheal mechanical ventilation) and neurological conditions including meningitis, head injury and cerebral infections.

The total sample consisted of participants with available MRI scan who had been assessed between February 2012 and December 2014. Participants gave full informed written consent and the study was approved by the appropriate local ethics committees, and in compliance with national legislation and the code of ethical principles for Medical Research Involving Human Subjects of the World Medical Association (Declaration of Helsinki).

### Socio-demographic, neuropsychological and neonatal data

Socio-economic status (SES) was evaluated in accordance with Her Majesty’s Stationary Office Occupational Classification criteria (46), using participants’ occupation at current assessment and parents’ jobs at birth. Categories were collapsed into two groups: high SES, which comprised managerial and professional roles, and low SES, which encompassed all the other occupations, including students and those who were currently unemployed. Hand preference was assessed using the Annett’s questionnaire (47). Participants were classified as left- or right-dominant when consistent answers were given to all items, or mixed when inconsistent responses were reported.

Intelligence Quotient (IQ) was assessed using the Wechsler Abbreviated Scale of Intelligence (WASI) (48) that includes two verbal tasks (vocabulary and similarities) and two visuo-spatial tasks (block design and matrix reasoning) to provide estimates of verbal, performance and full-scale IQ expressed as standard scores (*M* = 100, *SD* = 15). Psychiatric symptomatology was assessed with the CAARMS (35), a semi-structured interview that assess positive and negative symptoms, cognitive problems, emotional disturbance, behavioral changes, motor/physical changes and general psychopathology [see Kroll *et al.* (36) for further details]. The total psychopathology score, obtained as the sum of all items, was used as a measure of the non-specific psychiatric symptomatology found in our cohort (36).

For the preterm group only, neonatal variables included gestational age, birth weight, and severity of perinatal brain injury based on cranial ultrasound scan and classified as: 1) normal scan, 2) uncomplicated periventricular hemorrhage, or 3) periventricular hemorrhage and ventricular dilatation (49).

### MRI image acquisition and preprocessing

Acquisition parameters of structural MRI data are described in Supplement 1. FreeSurfer v5.1.0 (http://surfer.nmr.mgh.harvard.edu/) was used to carry out the automated pre-processing including volumetric segmentation and cortical surface reconstruction (50, 51); these were briefly described in Supplement 1. From a total of 166 participants with available T1 sequences, six (two controls) were excluded from the analyses because of failure of the automated image processing due to substantial topological defects, skull stripping problems or poor intensity segmentation. The total sample included 79 very preterm individuals and 81 controls.

### Local Gyrification Index computation

LGI was computed using the method proposed by Schaer *et al*. (38) and implemented in FreeSurfer (52). This involved three steps. First, an outer hull that tightly warps the pial cortical surface was created through a morphological closing algorithm. Second, hundreds of overlapping circular regions of interest were delineated over the outer surface and matched to the corresponding patches on the pial surface. At each vertex of the hull surface, LGI was calculated as the ratio between the perimeters of the two corresponding regions, which reflected the amount of cortex buried within the sulcal folds of each region. Finally, LGI values were propagated from the outer mesh to the pial mesh in order to obtain individual cortical maps, which were subsequently used for statistical comparisons.

Visual inspection of the automatically reconstructed surfaces revealed image defects for 14 participants (three controls), mainly attributable to poor signal intensity in the occipital lobe requiring manual pial edits. A sensitivity analysis showed that results with inclusion and exclusion of those participants were unchanged (Supplemental Figure S1), therefore here we report the main analyses results including the unedited scans. Additional information about quality check and parameter choices for LGI computation are reported in Supplement 1.

### Statistical analyses

Non-imaging data were analyzed using SPSS® Statistics 24.0 (53). Visual inspection plus skewness and kurtosis assessment were used to check for normal distribution of continuous variables. Parametric or non-parametric tests were used as appropriate.

To ascertain participation bias within the very preterm cohort, current study participants were compared to non-participants in terms of gestational age, birth weight, sex and neonatal US classification using independent-sample and chi-square tests respectively.

Differences in sex, handedness and SES between very preterm individuals and controls were assessed using Pearson Chi-square test (χ^2^), with Fisher’s exact correction if required. Group differences in white matter and intracranial volume, age at assessment and neuropsychological tests were analyzed using independent-samples *t*-tests or Mann-Whitney *U*-tests as appropriate.

Between-group differences in cortical gyrification were conducted at each vertex of the cortical surface in the left and right hemispheres, using individual maps smoothed with a 10-mm full-width half-maximum Gaussian kernel. Intracranial volume was used as covariate since decreased brain volume has previously been noted in this cohort at adolescence assessment in comparison with age-matched term-born controls (54). No difference was found in our study in intracranial volume (preterm: *M* [*SD*] = 1,458.76 [231.04] cm^3^; controls: *M* [*SD*] = 1,503.53 [205.15] cm^3^; *t*_158_ = 1.30, *p* = 0.197) and its relationship with LGI did not differ between the two groups.

First, statistical analysis on LGI across the entire mantle was conducted using ANCOVA with group (very preterm and control) and sex (male and female) as between-subject factors and intracranial volume as group-centered global covariate. Main effects of group and group-sex interactions were tested. Then, group differences were tested using a per-vertex regression model where individual cortical thickness maps were entered as per-vertex regressor, while covarying for global intracranial volume. The effect of sex was tested by repeating separate analyses for male and female participants. False Discovery Rate (FDR) correction *q* = 0.05 was applied to correct for multiple comparisons. Further, within-group analyses were conducted to investigate the vertex-wise correlations between LGI and: 1) full-scale IQ and 2) CAARMS measures in both very preterm and control participants, using an uncorrected significance threshold of *p* = 0.001. Group differences in the relationship between LGI and functional outcomes were also tested.

## Results

### Sample characteristics and neuropsychological assessment

Compared to the very preterm individuals who were not assessed at the follow-up at 30 years, the current participants had younger gestational age (*t*_375_ = 2.14, *p* = 0.033) and lighter birth weight (*t*_375_ = 2.05, *p* = 0.041). A higher proportion of males than females was assessed (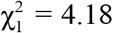, *p* = 0.041), and there were more individuals with complicated periventricular hemorrhage within the current sample (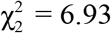, *p* = 0.031).

Table 1 displays socio-demographic and neonatal characteristics of the study sample. The two study groups did not differ in terms of sex, hand preference, parental SES at birth and participant SES at testing. Controls were younger than very preterm participants. At the neuropsychological assessment, very preterm participants had lower full-scale IQ than controls. There were no significant between-group differences in CAARMS total psychopathology. Additional measures of psychosocial functioning and mental health are reported in the Supplemental Table S1.

**Table 1.** Socio-demographic and neonatal characteristics, and results of the neuropsychological assessment of the very preterm and control groups. Descriptive statistics are given in percentages or mean (standard deviation), unless otherwise stated.

|  | Preterm<br>(*n* = 79) | Data<br>available (*n*) | Controls<br>(*n* = 81) | Data<br>available (*n*) | Test statistic | *p* value |
| --- | --- | --- | --- | --- | --- | --- |
| Sex (male/female) | 61/39 | 79 | 48/52 | 81 | $\chi^2_1 = 2.56$ | 0.109 |
| Age at assessment (in years) <sup>a</sup> | 30.24 (3.95) | 78 | 29.29 (5.02) | 79 | $U = 2,439.00$ | 0.024 |
| Handedness (right/left/mixed) | 83/10/7 | 60 | 88/7/5 | 57 | Fisher's exact $\chi^2_1 = 0.54$ | 0.856 |
| Subject SES at testing (low/high) | 53/47 | 78 | 51/49 | 80 | $\chi^2_1 = 0.03$ | 0.869 |
| Parental SES at birth (low/high) | 49/51 | 73 | 35/65 | 65 | $\chi^2_1 = 2.73$ | 0.099 |
| Neonatal characteristics |  |  |  |  |  |  |
| Birth weight (in grams) | 1280.82 (339.88) | 78 | n/a |  | – | – |
| Gestational age (in weeks) | 29.10 (2.21) | 78 | n/a |  | – | – |
| US classification (no-PVH/PVH/PVH+DIL) | 47/23/30 | 79 | n/a |  | – | – |
| Neuropsychological assessment |  |  |  |  |  |  |
| Full-scale IQ <sup>a</sup> | 107 (17) | 68 | 115 (14) | 64 | $U = 1,469.00$ | 0.001 |
| CAARMS total psychopathology <sup>a</sup> | 7 (14) | 65 | 4 (10) | 68 | $U = 1,847.50$ | 0.101 |

Note: missing data were mainly due to logistic problems at the beginning of the project and incomplete items preventing the computation of composite scores.
<sup>a</sup> Median (interquartile range).
CAARMS = Comprehensive Assessment of At-Risk Mental States. IQ = Intelligence Quotient. SES = socio-economic status.
Ultrasound Scan (US) classification: no-PVH = normal neonatal cranial ultrasound; PVH = uncomplicated periventricular hemorrhage; PVH + DIL = periventricular hemorrhage with ventricular dilatation.

### Gyrification differences between very preterm and control participants

Figure 1A shows the results of the ANCOVA testing for differences in LGI between very preterm individuals and controls, accounting for sex, while covarying for intracranial volume. Compared to term-born controls, very preterm participants showed reduced gyrification in two clusters, one in the left hemisphere (size = 61,036 mm^2^) and one in the right hemisphere (size = 56,036 mm^2^), both with peak in the insula (*p* < .001). The clusters extended to large portions of prefrontal, anterior temporal, medial and dorsal parietal, and occipital lobes bilaterally as well as in the left superior frontal gyrus. Moreover, preterm individuals showed increased gyrification in a small medial portion of the right temporal pole (size = 76 mm^2^, *p* < .01). There was no significant interaction between sex and group (*p* > 0.05).

**Figure 1.**
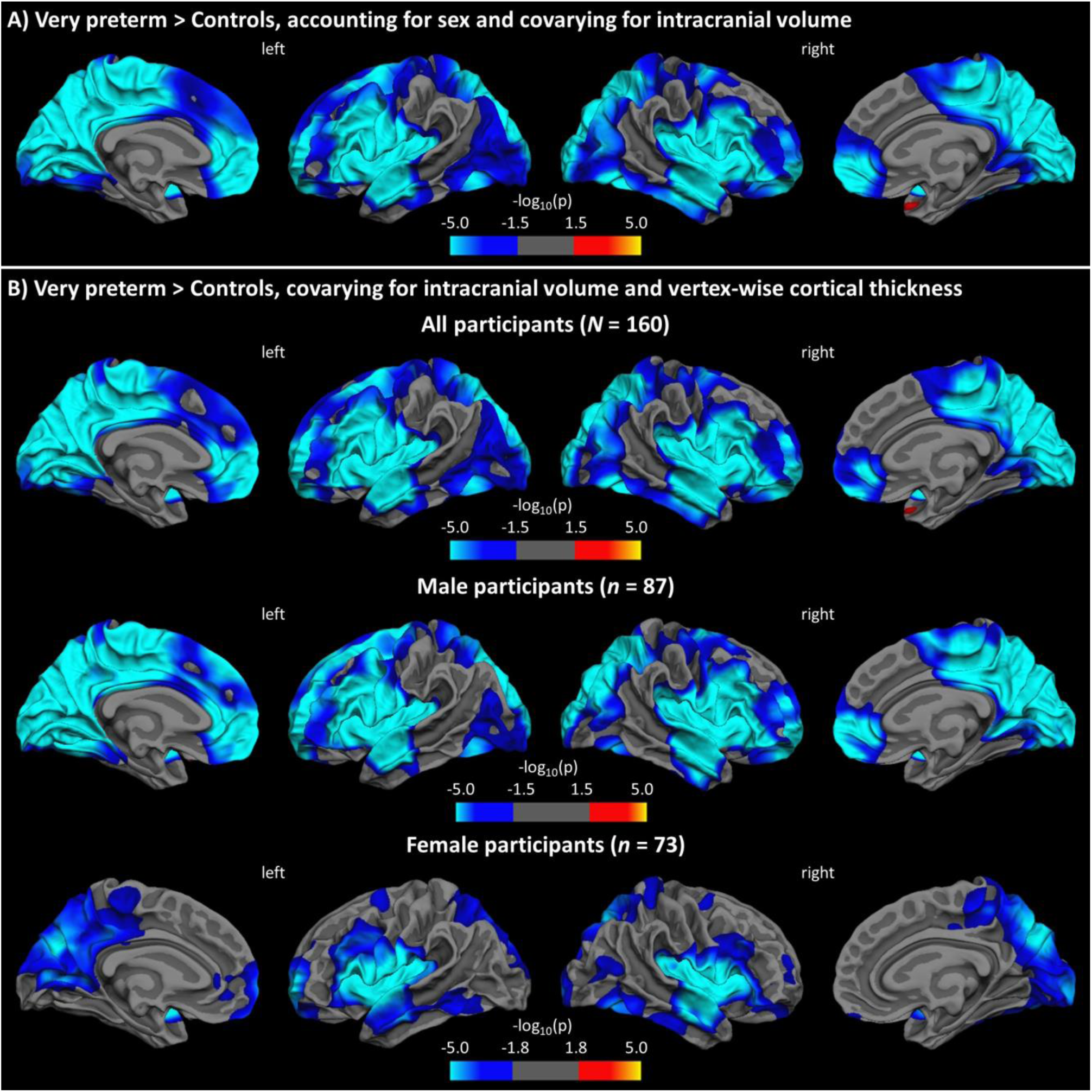
Results from general linear models testing for group differences in LGI: A) accounting for sex, with intracranial volume as global covariate; B) covarying for intracranial volume and vertex-wise cortical thickness for all (top row), male (middle row), and female (bottom row) participants. Statistical maps are displayed on the white surface, with FDR-corrected level of significance at 0.05.

Figure 1B and Table 2 show the results of the per-vertex regression model assessing group differences in LGI using intracranial volume and cortical thickness at each vertex of the surface as confounders. Very preterm participants showed significantly reduced cortical folding bilaterally in the insula, middle frontal and orbitofrontal gyri, anterior temporal lobe, prefrontal cortex and extended parts of the medial occipito-parietal cortices compared to term-born controls (see also Supplemental Figure S3). Separate analyses for sex yielded a similar pattern of hypogyria involving the insula, anterior temporal and medial parietal-occipital areas, whereas reduced folding in the frontal and lateral occipito-parietal lobes appeared more prominent in males than females. Conversely, the hypergyric region in the temporal lobe did not remain significant for either sex.

**Table 2.** Significant clusters showing group differences in LGI while covarying for intracranial volume and adjusting for vertex-wise cortical thickness (FDR correction *q* = 0.05).

| Subjects | Cortical region (H) | Cluster size,<br>mm<sup>2</sup> | $p$ value | Talaraich coordinates<br>of the max | | |
| --- | --- | --- | --- | --- | --- | --- |
|  |  |  |  | x | y | z |
| Total sample | Precentral (L) | 59,410 | < .001 | -39 | 0 | 15 |
|  | Precentral (R) | 54,116 | < .001 | 53 | -2 | 10 |
|  | Superior temporal (R) | 49 | < .05 | 40 | 11 | -26 |
| Males | Superior temporal (L) | 55,708 | < .001 | -43 | -20 | -6 |
|  | Lateral orbitofrontal (L) | 23 | < .05 | -26 | 18 | -19 |
|  | Superior temporal (R) | 23,094 | < .001 | 52 | -10 | -1 |
|  | Precuneus (R) | 29,031 | < .001 | 10 | -66 | 39 |
|  | Caudal middle frontal (R) | 3 | < .05 | 29 | 12 | 43 |
| Females | Precentral (L) | 14,684 | < .001 | -48 | -1 | 8 |
|  | Rostral middle frontal (L) | 2,471 | < .001 | -16 | 61 | -5 |
|  | Precuneus (L) | 12,472 | < .001 | -19 | -69 | 30 |
|  | Superior frontal (L) | 292 | < .01 | -19 | 5 | 56 |
|  | Medial orbitofrontal (L) | 110 | < .01 | -12 | 40 | -7 |
|  | Rostral anterior cingulate (L) | 62 | < .01 | -6 | 35 | 3 |
|  | Inferior parietal (L) | 73 | < .05 | -43 | -56 | 41 |
|  | Posterior cingulate (L) | 16 | < .05 | -8 | -24 | 29 |
|  | Precentral (L) | 19 | < .05 | -21 | -22 | 52 |
|  | Insula (R) | 28,447 | < .001 | 37 | -3 | 16 |
|  | Superior frontal (R) | 153 | < .01 | 22 | 1 | 53 |
|  | Inferior parietal (R) | 726 | < .01 | 44 | -71 | 18 |
|  | Rostral middle frontal (R) | 279 | < .01 | 22 | 55 | 15 |
|  | Rostral middle frontal (R) | 3 | < .05 | 39 | 42 | -1 |

H = hemisphere; L = left; R = right.

Overlapping results were obtained when intracranial volume was not included as a global covariate, with significant group effects found in nearly all the same regions (Supplemental Figure S2).

### Per-vertex correlations between cortical gyrification and full-scale IQ

Positive vertex-wise associations were found between LGI and full-scale IQ in separate-group analyses for preterm and control participants. In very preterm participants, associations between LGI and full-scale IQ were found in the left fusiform and lateral orbitofrontal cortex as well as in the right superior parietal gyrus (Figure 2A, Table 3). In control participants associations between LGI and full-scale IQ were found mainly in the right inferior parietal lobe, postcentral gyrus, lateral orbitofrontal and caudal middle frontal cortex, but also in the left pars orbitalis (Figure 2B; Table 3). There was no group-by-IQ interaction on LGI (*p* > 0.05).

**Table 3.** Significant associations of LGI with full-scale IQ (positive) and CAARMS total psychopathology (negative) for very preterm and control participants (all uncorrected *p* < .001).

| Measure | Subgroups | Cortical region (H) | Cluster size, mm<sup>2</sup> | Talaraich coordinates of the max |  |  |
| --- | --- | --- | --- | --- | --- | --- |
|  |  |  |  | x | y | z |
| Full-scale IQ | Very preterm | Lateral orbitofrontal (L) | 341 | -35 | 22 | -16 |
|  |  | Fusiform (L) | 614 | -29 | -74 | -6 |
|  |  | Superior parietal (R) | 94 | 31 | -48 | 61 |
|  | Controls | Pars orbitalis (L) | 45 | -40 | 29 | -13 |
|  |  | Inferior parietal (R) | 1,486 | 45 | -61 | 26 |
|  |  | Postcentral (R) | 247 | 25 | -36 | 50 |
|  |  | Superior frontal (R) | 303 | 17 | 20 | 52 |
|  |  | Lateral orbitofrontal (R) | 676 | 36 | 29 | -5 |
|  |  | Caudal middle frontal (R) | 235 | 36 | 7 | 34 |
|  |  | Precentral (R) | 24 | 25 | -10 | 60 |
| CAARMS | Very preterm | Precentral (L) | 48 | -17 | -19 | 67 |
|  |  | Postcentral (L) | 14 | -34 | -31 | 57 |
|  |  | Precentral (L) | 10 | -32 | -9 | 49 |
|  |  | Paracentral (R) | 123 | 10 | -29 | 49 |
|  |  | Postcentral (R) | 26 | 30 | -31 | 58 |
|  |  | Insula (R) | 69 | 37 | 0 | -6 |
|  |  | Insula (R) | 31 | 31 | 18 | -4 |
|  | Control | Supramarginal (R) | 184 | 49 | -25 | 26 |

H = hemisphere; L = left; R = right.

**Figure 2.**
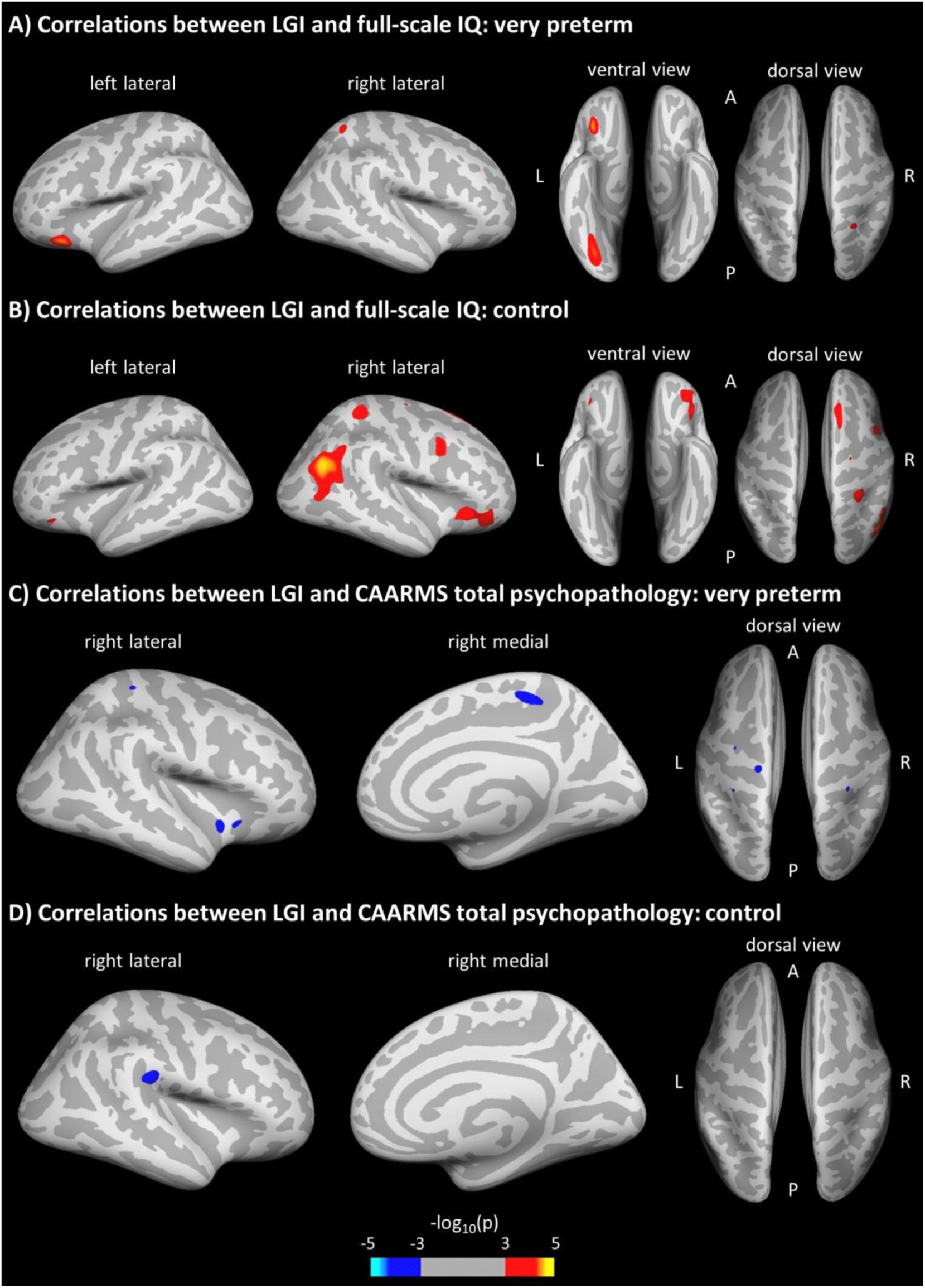
Vertex-wise associations of LGI with full-scale IQ and CAARMS total psychopathology for very preterm and control participants. Statistical maps are displayed on the inflated surface, with uncorrected level of significance *p* = 0.001. LGI = local gyrification index.

### Per-vertex correlations between cortical gyrification and CAARMS total psychopathology

Vertex-wise correlations between LGI and CAARMS total psychopathology scores were conducted after removal of two participants (one very preterm and one control) who were identified as outliers with unusually high CAARMS scores. Negative correlations were found throughout, but these involved different brain areas in each group. Very preterm participants displayed significant associations between LGI and CAARMS total psychopathology scores in the left precentral, bilateral postcentral, right paracentral regions as well as in the right insula (Figure 2C, Table 3). Control participants showed only a significant association in the right supramarginal gyrus (Figure 2D, Table 3). There was no group-by-CAARMS interaction on LGI (*p* > 0.05).

## Discussion

Using a three-dimensional measure of gyrification, this study showed altered gyrification in a large portion of the cortical mantle in adults who were born very preterm compared to age-matched term-born controls. Specifically, our results confirm the primary hypothesis of mostly reduced cortical folding, in line with previous evidence involving the orbitofrontal (26), insular and superior temporal lobes (20, 23) in preterm children and adolescents. Our observations partially confirm the results by Hedderich *et al*. (28), who reported widespread curvature reduction in very preterm/low-birth-weight adults extended to the parietal and temporal lobes, but not to the medial and lateral superior frontal cortex. Differences between the two studies might arise from the different 3D surface-based proxies of folding complexity.

In addition, the current study found a small hypergyria in the temporal lobe, but this pattern should be interpreted cautiously due to its inconsistency, as demonstrated in the sex-separate comparisons. Kesler et al. (24) demonstrated increased temporal lobe folding using a global 2D gyrification index in very preterm children, but other studies reported reduced folding complexity in this lobe (18, 23). The current study with a regional proxy suggests a specific vulnerability to both hypo- and hyper-gyrification of different temporal regions, although the underpinning mechanism remains to be elucidated (23). In line with the previous literature, our results altogether demonstrate significant gyrification abnormalities (both increased and decreased) in very preterm subjects as compared to term-born controls.

Sex did not show significant effects on gyrification, possibly because adjustment for brain volume accounted for folding differences between male and female adults (55). Nonetheless, sex-specific comparisons suggested that folding alterations were more prominent in male adults, in line with extensive research showing higher vulnerability for male preterm infants to neurodevelopmental morbidities (56–58) as well as structural (59) and functional (60) brain alterations. In addition, preterm girls have a more “compact” brain, with higher degree of cortical folding but smaller brain volume (61), which may have important implications for later outcomes and psychopathology (62). Furthermore, vertex-wise adjustment for cortical thickness, which is an indicator of brain maturation during childhood and adolescence (63), did not account for differences in adulthood on gyrification, which instead reflects early brain development during fetal life (37). Therefore, reduced gyrification detected in our adult sample could be interpreted as the result of an early alteration to the folding process in perinatal life that persists beyond cortical maturation programmed to occur later in life.

Using a whole brain approach, we found the strongest gyrification reduction in the very preterm sample bilaterally in the insula, a deeply buried region characterized by elevated convolution complexity across lifespan (12, 64). The insula, the first lobe to differentiate in fetal life, starts sulcation/gyrification at 16 weeks of gestation (4, 65) and undergoes critical development between 21 and 32 weeks (66), making it particularly vulnerable to alterations following very preterm birth. Nonetheless, the hypogyria consistently extended beyond the insula to the anterior temporal lobe, inferior frontal lobe, orbitofrontal cortex, and medial occipito-parietal regions, and its robustness was demonstrated in both male and female participants. Our observation shows a striking overlap with the results of a whole-brain study in individuals with adult-onset schizophrenia reporting reduced gyrification in the insular region with extensions to the inferior frontal gyrus and superior temporal gyrus (30). Both increased and decreased gyrification in these areas were also found in another sample of adolescents with schizophrenia (67), as well as in other psychotic and nonpsychotic conditions, such as bipolar disorder and depression (32), which are more prevalent in the preterm population (3). Therefore, our findings imply that aberrant gyrification seen in individuals with various psychiatric disorders could at least partly be explained by neurodevelopmental disruptions occurring very early in life.

Gyrification showed a positive linear relationship with full-scale IQ in both very preterm and control participants, especially in the left orbitofrontal and right superior parietal regions. A distinctive association for the preterm group was found in the left fusiform. For the control group, unique associations involved the middle and inferior frontal lobe as well as the temporo-parietal junction in the right hemisphere. These findings corroborate evidence of a wide fronto-parietal network sustaining the association between WASI full-scale IQ and LGI in a large cohort of healthy adults (40). Notably, in the current study the significant group difference in full-scale IQ probably reflects not so much an impairment of very preterm participants but rather an overperformance of control participants who tend to score +1 SD above the test mean, arguably because many were university students (Supplement, Table S1). Therefore, it is plausible that the network containment and the reduced involvement of associative cortices might be implicated in the lowered IQ found in very preterm participants in comparison to term-born controls.

Finally, exploratory analyses showed the same direction of negative associations between local gyrification and psychopathology scores in both very preterm and control groups, but different spatial distribution across the whole brain. In controls, one significant association was found only in the right supramarginal gyrus, an area specifically implicated in empathic judgements (68) and social cognition through the frontoparietal mirror system (69). Instead, in the very preterm group the association involved the central and paracentral lobules, corresponding to the primary sensory-motor cortices (70), and the right insula, specialized in interoception, multimodal and somatosensorial processing (71, 72) and emotional awareness (73). The CAARMS, a tool specifically designated to recognize the subclinical symptomatology of a first psychotic episode (35), includes questions about perception and body-sensation abnormalities as well as emotional-affective dysfunction. Therefore, while higher gyrification is associated with better adult mental health in both preterm and term-born individuals, the regional specificity of the relationships suggests that the link between early gyrification process and later psychiatric problems is qualitatively different in the two groups because of the different neural substrates. The current study is the first to show significant associations between gyrification and psychiatric outcomes in preterm individuals, which were not examined (24, 26, 28) or demonstrated (27) previously. While the paucity of research and methodological differences might explain this novel finding, it is also possible that such associations arise only later in adult life, in continuity with childhood psychopathology (74) even when considering subclinical symptoms (75). The initial delay of the folding process following very preterm birth (17–20) might contribute to the faster LGI decline seen in various psychiatric diagnoses (64). However, more research is necessary in future to elucidate the nature of the relationship between morphological features and dimensional functional outcomes, i.e. to establish concurrent and predictive associations throughout life and in relation to sex differences (62).

Several limitations should be considered when evaluating this research. Attrition analysis suggests that we considered a high-risk sample because all perinatal risk factors were not favourable in our preterm group in comparison to the non-returning cohort, unlike what commonly occur in longitudinal studies (76). However, the neuroimaging nature of this study might have involved high-functioning participants who were able to undergo an MRI scan (77), as suggested also by the similar psychopathology levels reported in preterm and control participants. Furthermore, our preterm cohort born three decades ago may not be representative of more recent preterm cohorts who receive novel neuroprotective interventions (78), but long-term sequelae seem to persist in spite of advancements in neonatal care (79). Our control group had a slightly younger mean age at assessment compared to the preterm group; data plotting also showed that controls comprised a few older individuals. However, since LGI decreases with age in healthy individuals (14, 64), the presence of these participants is expected to have attenuated group differences in our study. Finally, another limitation regards the use of the CAARMS, which is designed for individuals at ultra-high risk of psychosis. Future research should employ a diagnostic interview that is for use with the general population and examines various psychiatric disorders (80), or self-reported psychopathology assessment, that might be sensitive to subclinical symptoms (81).

To conclude, the current study found widespread reduced cortical gyrification in very preterm adults compared to term-born controls, particularly in the middle and inferior frontal, superior temporal and medial occipito-parietal regions. It also described specific regional associations of local gyrification with cognitive ability and mental health outcomes in adulthood. Overall, this study demonstrates that gyrification measured in adult life provides a fingerprint for very preterm birth, with wide aberrations persisting in adult life and possibly implicated in functional outcomes.

## Acknowledgments

The study was funded by a Medical Research Council, UK research grant (ref. MR/K004867/1) to CN. LP acknowledges support from the Tanna Schulich Chair of Neuroscience and Mental Health and the Opportunities Fund of the Academic Health Sciences Centre Alternative Funding Plan of the Academic Medical Organization of Southwestern Ontario (AMOSO).

We thank our study participants for their continuing help. We also thank the National Institute for Health Research (NIHR) Biomedical Research Center at South London and Maudsley NHS Foundation Trust and King’s College London for supporting the neuroimaging facilities used in our study.

