## Supplement 1 for "Altered cortical gyrification in adults who were born very preterm and its associations with cognition and mental health"

### Supplemental Methods & Material

***MRI image acquisition***

Structural MRI data were acquired on a SIGNA HDx 3.0 Tesla MR scanner (GE Healthcare, USA). T1-weighted images were collected using an Enhanced Fast Gradient Echo 3-Dimensional (efgre3D) sequence with the following parameters: repetition time (TR) = 7.1 ms, echo time (TE) = 2.8 ms, inversion time (TI) = 450 ms, flip angle = 20°, field of view (FOV) = 280 mm, matrix = 256 x 256. Each volume contained 196 contiguous slices (slice thickness = 1.1 mm) with reconstructed voxel resolution of 1.1 mm^3^.

***MRI image preprocessing***

Firstly, the volume-based stream images included motion correction, transformation to the Talairach space, intensity normalization, removal of non-brain tissues, volumetric subcortical labelling and white matter segmentation. This pipeline produced volumetric estimates of intracranial space, white and grey matter tissues as well as various subcortical structures for each subject.

Secondly, the surface-based stream used the white matter volume to generate hemispheric surface maps, which were modelled as a mesh of triangular faces defined by ~150,000 vertices. These surface maps were then deformed to represent the white matter surface (grey-white boundary) and the pial surface (grey-CSF boundary) based on intensity gradients and neighborhood constraints. After inflation and spherical registration, a spherical atlas was mapped back onto the average inter-subject representation and then on the individual inflated surfaces with fine tuning with tissue boundaries. Individual cortical thickness maps were obtained by calculating the distance between two corresponding vertices on the white and pial surfaces.

***Quality check and parameter choices for LGI computation***

We manually checked all reconstructions to ensure that the ‘potholes’ and ‘handles’ were identified on the reconstructed grey matter surface. In particular, for the areas susceptible to MR inhomogeneities (temporal pole, inferior temporal surface, medial orbital region), we carefully evaluated the automated segmentations to ensure the accuracy of our results.

For computing LGI, the radius of the circular region of included surface (‘patch’) was set to 25 mm default in line with Schaer et al. ^[1]^, and multiple clinical studies that demonstrated sensitivity of this radius subsequently (2-6). This radius ensures that more than one sulcus is included in each calculation (thus capturing at least one ‘ridge’ and one ‘valley’) without diluting the local maxima needed to demonstrate peak vertex effects. Notably, the effect of statistical smoothing is negligible as the cortical LGI maps are already smoothed by the choice of 25 mm radius. Nevertheless, in keeping with our prior studies, we used 10 mm smoothing, recommended by Schaer et al.^1^.

We used ICV as global covariate, but the exclusion of this covariate had negligible effects on the final results, as shown in Supplemental Figure S2.

1. 1. Schaer M, Cuadra MB, Tamarit L, Lazeyras F, Eliez S, Thiran J-P (2008): A surface-based approach to quantify local cortical gyrification. *IEEE Trans Med Imaging*. 27: 161–170.
