## Supplementary material for "Altered cortical gyrification in adults who were born very preterm and its associations with cognition and mental health": Table S1

### Further assessment of psychosocial functioning and mental health

In terms of SES, there was a different distribution of very preterm and control subjects across the socio-economic categories, due to a larger proportion of students in the control sample at a Bonferroni-corrected level of significance (0.05/7 = 0.007).

Participants’ level of functioning was measured with the total score on the Role Functioning Scale^[1]^, a clinical interview that assesses the domains of working productivity, independent living and self-care, immediate social network relationship and extended social network. Two measures of participants’ mental health at testing were also collected: 1) the total score on the 12-item General Health Questionnaire (GHQ-12)^[2]^ to evaluate anxiety and depression symptoms, and 2) the total score on the 21-item Peters’ Delusional Inventory (PDI-21)^[3]^ to assess psychosis tendency. Finally, autistic traits were assessed using the Autism-Spectrum Quotient (AQ-10)^[4]^.

Results of the neuropsychological assessment on these additional measures are reported in Table S1. There were no differences between preterm individuals and term-born controls on any of the outcomes. This further suggests that psychiatric problems, as reported in epidemiological studies, may be difficult to detect as full-blown disturbance among very preterm adults, especially in MRI studies.

| **Table S1.** Descriptive statistics (percentage, median and interquartile range) and between-group comparisons (Mann-Whitney *U*-test) of the additional of psychological and mental health assessment. | | | | | | |
| --- | --- | --- | --- | --- | --- | --- |
|  | Preterm (*n* = 79) | Data available (*n*) | Controls (*n* = 81) | Data available (*n*) | Test statistic | *p* value |
| Socio-economic categories: |  |  |  |  | χ = 19.1 | 0.002 |
| I (professional) | 8.8% | - | 4.9% | - |  |  |
| II (intermediate) | 37.9% | - | 43.2% | - |  |  |
| III (skilled manual and non-manual) | 29.1% | - | 25.9% | - |  |  |
| IV (semi-skilled manual) | 3.8% | - | 0% | - |  |  |
| V (unskilled manual) | 1.3% | - | 0% | - |  |  |
| Students | 1.3% | - | 17.3% | - |  |  |
| Unemployed/out of work | 16.5% | - | 7.4% | - |  |  |
| Missing | 1.3% | - | 1.3% | - |  |  |
| Role Functioning Scale | 26 (2) | 62 | 26 (2) | 60 | *U* = 1,595.0 | 0.160 |
| General Health Questionnaire | 10 (8) | 74 | 9 (6) | 71 | *U* = 2,511.0 | 0.645 |
| Peter’s Delusion Inventory | 20 (49) | 60 | 18 (36) | 63 | *U* = 1,740.5 | 0.448 |
| Autism Questionnaire 10 | 2 (2) | 41 | 2 (3) | 41 | *U* = 704.0 | 0.195 |

1. Goodman SH, Sewell DR, Cooley EL, Leavitt N (1993): Assessing levels of adaptive functioning: the Role Functioning Scale. *Community Ment Health J*. 29: 119–131.
2. Golderberg D, Williams P (1998): *A user’s guide to the General Health Questionnaire*. Windsor, UK: NFER-Nelson.
3. Peters ER, Joseph SA, Garety PA (1999): Measurement of delusional ideation in the normal population : introducting the PDI (Peters et al. Delusions Inventory). *Schizophr Bull*. 25: 553–576.
4. Allison C, Auyeung B, Baron-Cohen S (2012): Toward brief “red flags” for autism screening: The short Autism Spectrum Quotient and the short Quantitative Checklist in 1,000 cases and 3,000 controls. *J Am Acad Child Adolesc Psychiatry*. 51: 202-212.e7.
