## Supplementary material for "Altered cortical gyrification in adults who were born very preterm and its associations with cognition and mental health": Figure S1

### Sensitivity analyses

Sensitivity analyses that excluded subjects with defective reconstructed surfaces were conducted to examine the robustness of the main neuroimaging comparisons assessing between-group difference in gyrification across the entire mantle. Sensitivity analyses were conducted using scans from 68 very preterm participants and 78 controls.

Supplemental Figure S1 extensively overlap with Figure 1A in the main article, corroborating the results obtained when subjects with defective reconstructed surfaces are included in the analyses.

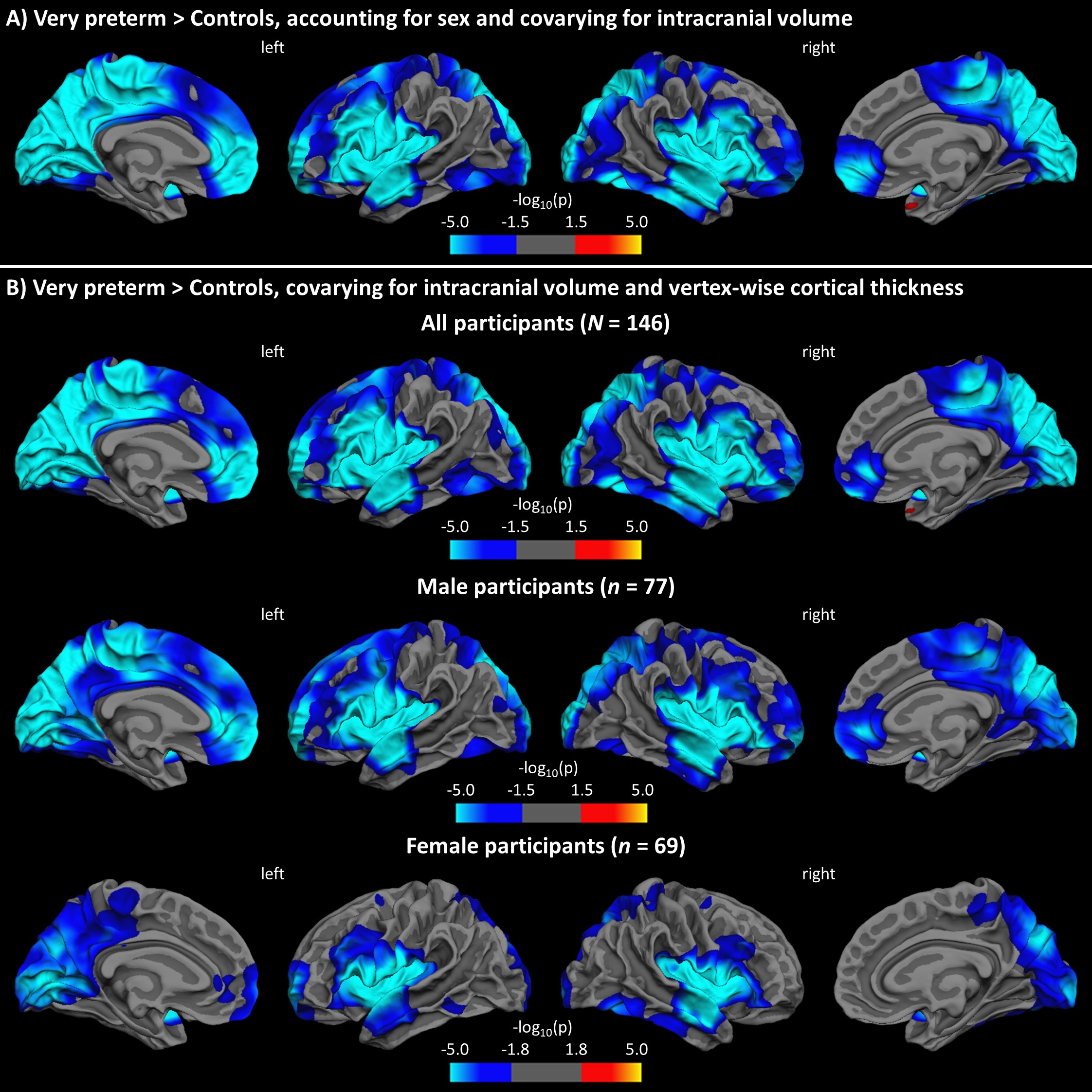

**Supplemental Figure S1.** White surface maps showing local patterns of hypogyria (cold colours) and hypergyria (warm colours) in very preterm participants compared to controls, after excluding subjects with defective reconstructed surface. A) Results of GLM accounting for sex and covarying for intracranial volume. B) Results of GLM covarying for intracranial volume and vertex-wise cortical thickness for all (top row), male (middle row), and female (bottom row) participants. FDR-corrected level of significance at *q* = 0.05

| **Supplemental Table S2**. Significant clusters showing group differences in local gyrification while covarying vertex-wise cortical thickness, after excluding subjects with defective reconstructed surface (FDR correction *q* = .05). | | | | | | |
| --- | --- | --- | --- | --- | --- | --- |
| Subjects | Cortical region (H) | Cluster size,  mm^2^ | *p* value | Talaraich coordinates  of the max | | |
|  |  |  |  | x | y | z |
| Total sample | Precentral (L) | 55,592 | < .001 | -54 | -1 | 7 |
|  | Precentral (R) | 50,611 | < .001 | 54 | -1 | 9 |
|  | Temporal pole (R) | 29 | < .01 | 40 | 11 | -26 |
| Males | Precentral (L) | 51,890 | < .001 | -55 | 0 | 6 |
|  | Precentral (R) | 47,899 | < .001 | 54 | -1 | 8 |
| Females | Superior temporal (L) | 12,390 | < .001 | -53 | -5 | -3 |
|  | Pericalcarine (L) | 13,604 | < .001 | -14 | -79 | 13 |
|  | Rostral middle frontal (L) | 2,113 | < .001 | -15 | 62 | -6 |
|  | Rostral anterior cingulate (L) | 286 | < .01 | -11 | 42 | 3 |
|  | Rostral anterior cingulate (L) | 74 | < .01 | -6 | 36 | 2 |
|  | Inferior temporal (L) | 325 | < .01 | -55 | -49 | -10 |
|  | Lateral orbitofrontal (L) | 36 | < .01 | -22 | 30 | -11 |
|  | Superior frontal (L) | 50 | < .05 | -19 | 5 | 56 |
|  | Posterior cingulate (L) | 4 | < .05 | -8 | -25 | -28 |
|  | Insula (R) | 25,265 | < .001 | 36 | -2 | 16 |
|  | Superior frontal (R) | 68 | < .01 | 22 | 1 | 54 |
| H = hemisphere; L = left; R = right. | | | | | | |
