## Supplementary material for "Altered cortical gyrification in adults who were born very preterm and its associations with cognition and mental health": Figure S2

### Unadjusted results without covarying for intracranial volume

The main neuroimaging comparisons assessing between-group difference in gyrification across the entire mantle were repeated without using intracranial volume as global covariate.

Supplemental Figure S2 extensively overlaps with Figure 1A in the main article, demonstrating marginal effects of intracranial volume correction on LGI group differences.

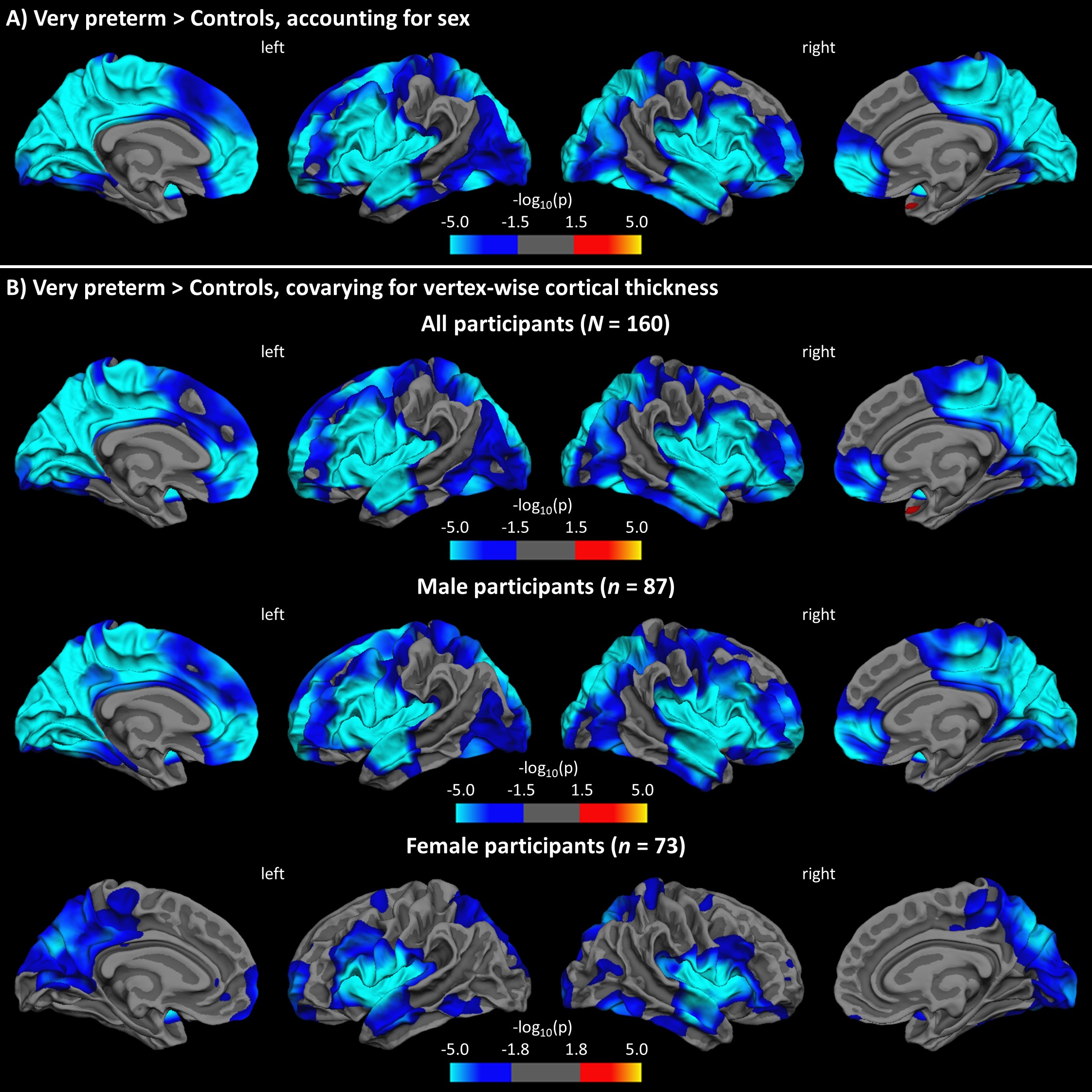

**Supplemental Figure S2.** White surface maps showing group differences in LGI: A) accounting for sex, and B) covarying for vertex-wise cortical thickness for all (top row), male (middle row) and female (bottom row) participants. Cold and warm colours indicate regiornal patterns of hypogyria and hypergyria respectively in very preterm participants compared to controls. FDR-corrected level of significance at 0.05.

| **Supplemental Table S3**. Significant clusters showing group differences in local gyrification while covarying vertex-wise cortical thickness (FDR correction at *q* = 0.05). | | | | | | |
| --- | --- | --- | --- | --- | --- | --- |
| Subjects | Cortical region (H) | Cluster size,  mm^2^ | *p* value | Talaraich coordinates of the max | | |
|  |  |  |  | x | y | z |
| Total sample | Precentral (L) | 58,746 | < .001 | -40 | 2 | 13 |
|  | Precentral (R) | 53,131 | < .001 | 53 | -2 | 10 |
|  | Superior temporal (R) | 60 | < .01 | 40 | 10 | -26 |
| Males | Superior temporal (L) | 57,962 | < .001 | -52 | -5 | -5 |
|  | Superior temporal (R) | 56,407 | < .001 | 52 | -9 | -1 |
|  | Temporal pole (R) | 13 | < .01 | 31 | 2 | -25 |
| Females | Precentral (L) | 12,626 | < .001 | -47 | -4 | 10 |
|  | Precuneus (L) | 11,806 | < .001 | -18 | -67 | 32 |
|  | Rostral middle frontal (L) | 2,009 | < .001 | -16 | 61 | -6 |
|  | Inferior temporal (L) | 1,099 | < .01 | -55 | -52 | -10 |
|  | Superior frontal (L) | 262 | < .01 | -19 | 6 | 57 |
|  | Lateral orbitofrontal (L) | 66 | < .01 | -22 | 30 | -11 |
|  | Medial orbitofrontal (L) | 49 | < .01 | -12 | 40 | -7 |
|  | Rostral middle frontal (L) | 47 | < .05 | -20 | 49 | 22 |
|  | Posterior cingulate (L) | 13 | < .05 | -8 | -25 | 29 |
|  | Rostral anterior cingulate (L) | 17 | < .05 | -6 | 36 | 2 |
|  | Inferior parietal (L) | 33 | < .05 | -43 | -56 | 41 |
|  | Insula (R) | 27,011 | < .001 | 37 | -2 | 16 |
|  | Superior frontal (R) | 130 | < .01 | 22 | 0 | 54 |
|  | Inferior parietal (R) | 590 | < .01 | 44 | -71 | 18 |
|  | Rostral middle frontal (R) | 48 | < .05 | 22 | 56 | 13 |
|  | Rostral middle frontal (R) | 20 | < .05 | 24 | 52 | 4 |
| H = hemisphere; L = left; R = right. | | | | | | |
