## Supplementary material for "Altered cortical gyrification in adults who were born very preterm and its associations with cognition and mental health": Figure S3

### Sex and preterm birth

We further assessed the effect of sex on gyrification within the preterm group by conducting a *t*-test comparison between male and female very preterm subjects. The uncorrected results (*p* < 0.01) are displayed in the Figure S3 below. None of these cluster survived FDR correction. This confirms that the lack of sex-by-group interaction may relate to not having sufficient sample size to seek such interactions in the current study.

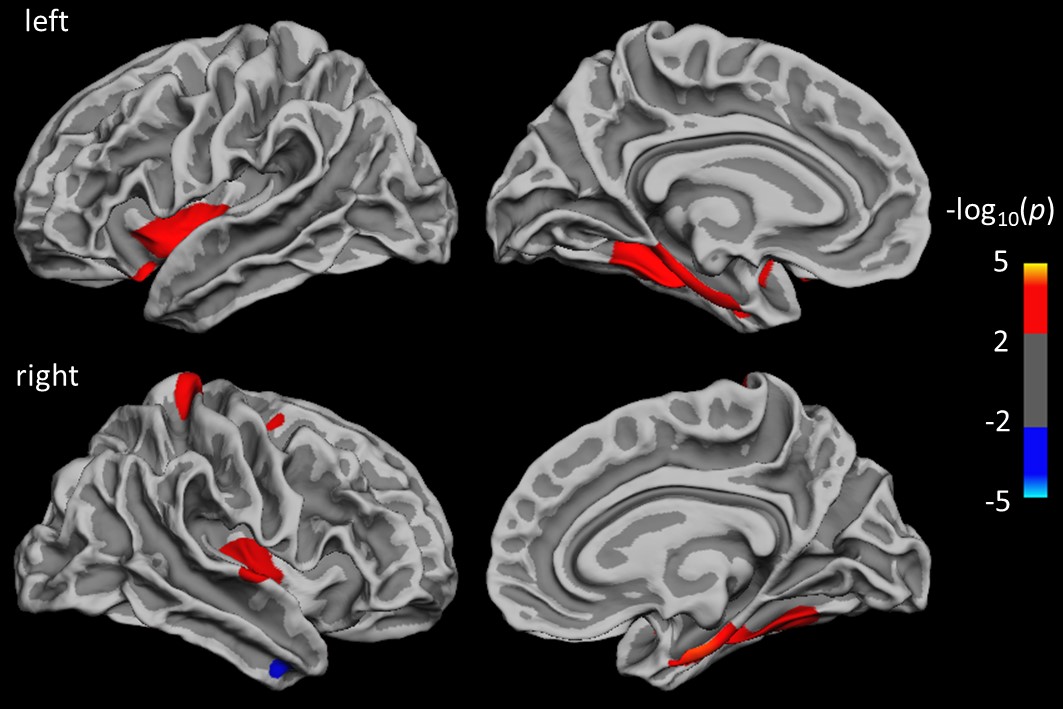

**Figure S3.** White surface maps showing LGI differences in male compared to female subjects within the very preterm group (uncorrected *p* < 0.01).
